# When indicator species identified from a network structure predict biodiversity

**DOI:** 10.64898/2026.02.02.703325

**Authors:** Ilhem Bouderbala, Daniel Fortin, Junior A. Tremblay, Christian Hébert, Antoine Allard, Patrick Desrosiers

## Abstract

Comprehensive biodiversity surveys are difficult to sustain across large and environmentally heterogeneous regions, creating a need for approaches that can recover community composition from a limited number of observed species. We ask whether a small set of indicator species selected from species co-occurrence networks can be used to reconstruct the occurrence of the remaining assemblage. We develop an integrated framework combining probabilistic co-occurrence networks, network-based indicator selection, and species distribution modeling, and evaluate it across contrasting bird and beetle assemblages prediction. Network-guided indicator selection supported species-level reconstruction across both taxa and was competitive with, and in several assemblages superior to, conventional selection strategies. Meaningful predictive performance was also retained for several low-prevalence species. However, indicator-based models generally matched or underperformed models based directly on climate, suggesting that part of the information carried by co-occurrence networks reflects environmental gradients already captured by measured climatic variables. The clearest gains beyond climate occurred in an assemblage characterized by a larger number of significant associations, broader spatial extent, and greater ecological heterogeneity, suggesting that the value of network-guided indicators may increase when richer association structure is available. Together, these results show both the potential and the limits of co-occurrence networks for identifying compact sets of species that can support species-level biodiversity reconstruction.

## 1 Introduction

Climate change and other anthropogenic pressures are altering global biodiversity, threatening ecosystem integrity and the services ecosystems provide to human societies (Thuiller et al., 2011; Pachauri et al., 2014; Zhang and Liang, 2014). Rising temperatures, shifting precipitation regimes, and more frequent extreme climatic events are driving species-range shifts, altering population dynamics, and reshaping species associations (Duveneck et al., 2014). For example, the northward expansion of thermophilous hardwoods at the expense of boreal conifers illustrates the spatial reorganization of ecological communities under climate pressure (Boulanger and Puigdevall, 2021). Land-use change, pollution, and biological invasions further accelerate biodiversity loss and ecological disruption (Masson-Delmotte et al., 2018; Labadie et al., 2021). These changes can affect ecosystem resilience and services such as pollination, pest control, and carbon sequestration (Chapin Iii et al., 2000; Oliver et al., 2015; Huang et al., 2021).

Accurate monitoring of how species assemblages reorganize across environmental gradients is therefore essential for ecological forecasting and conservation planning (Azeria et al., 2009; Dommain et al., 2020). Comprehensive biodiversity surveys, however, remain limited by logistical, taxonomic, and financial constraints, as well as imperfect detection that can generate false absences and biased community estimates (Kim and Byrne, 2006; MacKenzie et al., 2004). Indicator species (IS) offer a practical alternative: the presence or absence of a limited set of species may serve as a proxy for broader ecological conditions or biodiversity patterns (Dufrêne and Legendre, 1997; Chu et al., 2021; Terrigeol et al., 2022; Siddig et al., 2016). Yet surrogate effectiveness depends on the surrogate type, habitat, spatial scale, and analytical framework used for evaluation (Mellin et al., 2011). Indicator selection is itself consequential because it determines the ecological interpretation and potential utility of the resulting monitoring metric (Butler et al., 2012). An indicator should therefore be assessed not only by its association with aggregate biodiversity metrics, but also by whether it represents variation in community composition and provides predictive information beyond the environmental covariates available for monitoring.

Most indicator-species studies have focused on aggregate descriptors such as species richness or diversity indices rather than on reconstructing the occurrence patterns of the individual species that compose assemblages. Richness provides a useful summary of community structure, but it does not identify which species are present or absent and may therefore miss compositional reorganization across environmental gradients. Moving from richness prediction to species-level occurrence reconstruction is thus a conceptual and methodological shift for indicator-based biodiversity monitoring. This challenge is particularly important for uncommon and poorly sampled species. Such taxa may contribute disproportionately to ecosystem functioning and resilience, yet sparse occurrence data can make species-distribution models unstable and prone to overfitting (Jeliazkov et al., 2022; Miller-ter Kuile et al., 2025; Breiner et al., 2015). Reliable reconstruction consequently requires species-level evaluation, rather than inference from in-sample fit or from aggregate richness alone.

Species co-occurrence networks provide one possible basis for selecting indicator species. However, co-occurrence patterns should not be interpreted as direct evidence of ecological interactions. Significant associations may arise from shared environmental responses, dispersal, sampling processes, or indirect effects mediated by other species; indirect effects can even reverse the apparent sign of pairwise associations (Blanchet et al., 2020; Cazelles et al., 2016; Harris, 2016). Null-model, probabilistic co-occurrence, and network-based approaches nevertheless provide useful tools for describing statistical structure in community data (Gotelli, 2000; Veech, 2013; Rivest and Ebouele, 2020). Their ecological interpretation and predictive utility should be kept separate: an association may summarize assemblage structure without improving prediction of a target species once measured environmental covariates are available (Blanchet et al., 2020; Cazelles et al., 2016; Tikhonov et al., 2017; Warton et al., 2015). We therefore treat co-occurrence networks as statistical representations of assemblage-level association rather than maps of direct ecological interactions.

The predictive value of indicator sets may also be context dependent. Species pools, interaction strengths, climatic conditions, and detection processes vary across spatial and environmental gradients, potentially changing both indicator identity and performance (Santillán et al., 2018; Terrigeol et al., 2022; Araújo and Rozenfeld, 2014). Few studies have explicitly quantified how reconstruction accuracy and uncertainty propagate across broad gradients characterized by strong community turnover (Bouderbala et al., 2023a). Recent empirical evaluations further show that surrogate relationships can vary across space and environmental conditions, even when selected indicators co-occur with a substantial portion of the broader community (Lindenmayer et al., 2024; Brunk et al., 2025). Evaluating indicator performance in the ecological context in which monitoring will occur is therefore essential for developing transferable monitoring strategies.

Here, we develop a scalable network-based framework to reconstruct community composition from a limited set of indicator species. We combine probabilistic co-occurrence modeling (Veech, 2013) with graph-theoretic selection of indicator sets (Morueta-Holme et al., 2016), using positive degree, negative degree, betweenness, a betweenness–closeness criterion, and a consensus criterion integrating the four rankings. We benchmark these network-based approaches against richness-based, random, and Dufrêne–Legendre indicator-value (IndVal) selection (Dufrêne and Legendre, 1997; Terrigeol et al., 2022). We apply the framework to breeding-bird and beetle assemblages across climate-defined assemblages in Québec, Canada. We ask whether network-selected indicators can reconstruct species-level occurrences, whether they remain informative for low-prevalence species, and whether they provide information beyond measured climate. By jointly evaluating association polarity, network topology, and species prevalence, the framework identifies when compact indicator sets can support composition-resolved biodiversity monitoring and when measured climate already contains most of the available predictive signal.

Here, we develop a scalable network-based framework to reconstruct community composition from the presence–absence of a small set of indicator species (IS) (Fig. 1). The framework proceeds in four steps. First, within each climate-defined assemblage, we build a signed co-occurrence network using the probabilistic model of Veech (Veech, 2013), retaining significant positive and negative associations. Second, we select candidate IS sets using five graph-theoretic criteria (Morueta-Holme et al., 2016): positive degree, negative degree, betweenness centrality (Csardi and Nepusz, 2006), a betweenness–closeness criterion (BC), and a Consensus criterion integrating these four rankings. We benchmark them against richness-based (Terrigeol et al., 2022), random, and Dufrêne–Legendre indicator-value (IndVal) selection (Dufrêne and Legendre, 1997). Third, we model the occurrence of each non-indicator species with logistic regression using indicator species alone, climate alone, or both, and evaluate predictions exclusively on held-out data. Fourth, we compare reconstruction performance among selection methods and against the climate-only baseline. Applying this framework to breeding-bird and beetle assemblages in Québec, Canada, we ask whether network-selected indicators can reconstruct species-level occurrences, whether they remain informative for low-prevalence species, and whether they provide information beyond measured climate. By jointly evaluating association polarity, network topology, and species prevalence, we identify when compact indicator sets can support composition-resolved biodiversity monitoring and when measured climate already contains most of the available predictive signal.

**Figure 1.**
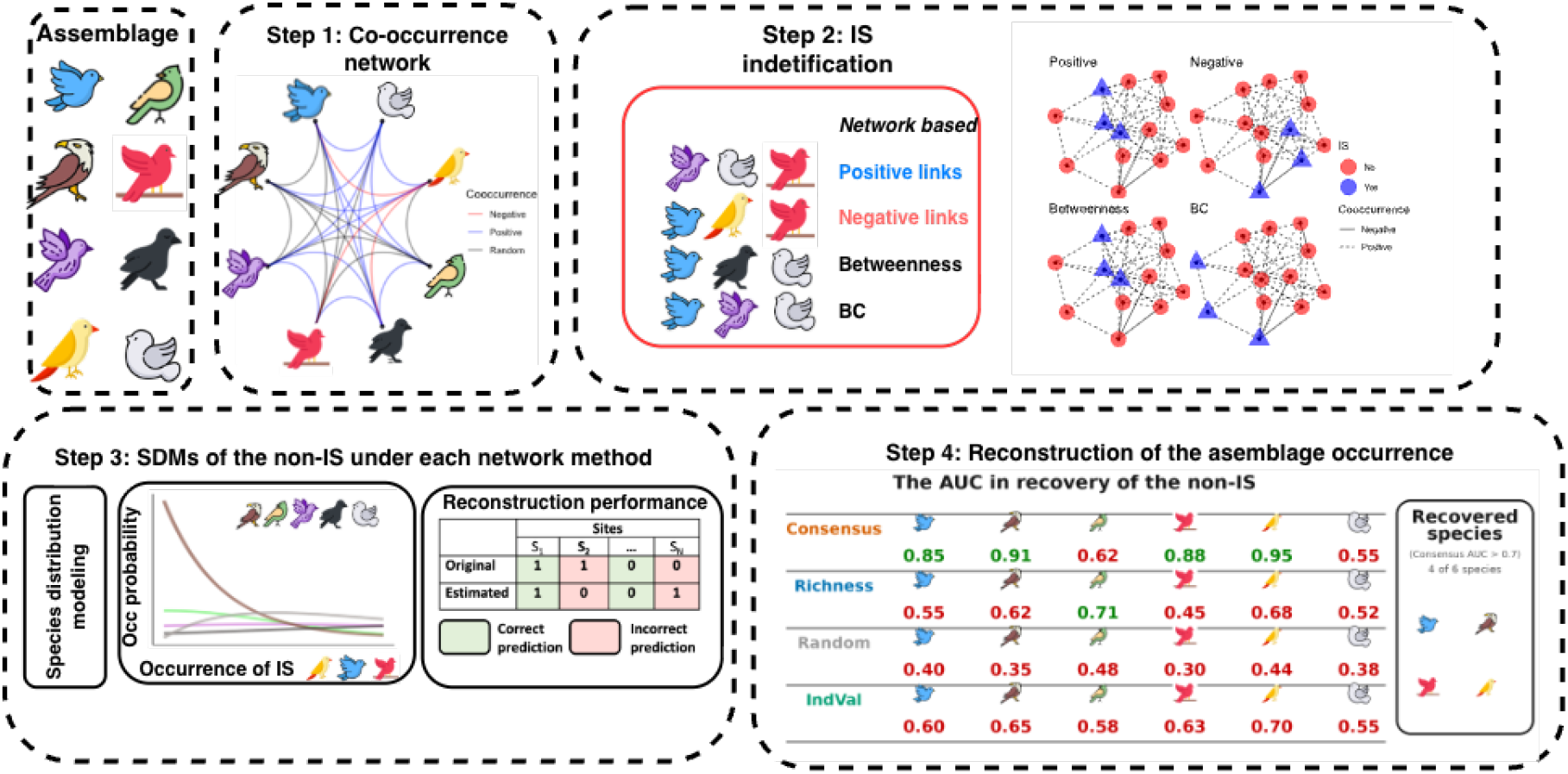
Conceptual framework for assemblage occurrence reconstruction. Step 1: signed co-occurrence networks are built within each climate-defined assemblage using Veech’s probabilistic model (Veech, 2013). Step 2: indicator species are selected using five network-based criteria (Positive, Negative, Betweenness, BC, and Consensus) and benchmarked against Richness, Random, and IndVal. Step 3: occurrence of non- indicator species is modeled using GLMs based on IS, Environment, or Environment+IS and evaluated by 5-fold cross-validation. Step 4: reconstruction performance is compared across selection methods and against the environment-only baseline.

## 2 Results

### 2.1 Species co-occurrence networks

The two taxa differed sharply in how their species were connected: birds formed small, dense networks, whereas beetles formed larger but sparser ones (Figs. S1–S2). Bird networks were compact and densely connected: the seven assemblages contained 34–55 species (74 to 457 sites), yet 6–22% of all possible species pairs were significantly associated, and positive associations dominated in every cluster. Beetle assemblages were two to three times richer (112–145 species across 42–121 sites) but produced sparser networks, with densities of only about 5–12%. Higher richness therefore did not translate into denser association structure. However, because the number of possible species pairs grows rapidly with richness, sparse beetle networks could still contain many associations in absolute terms. Beetles Cluster 3, the richest assemblage (145 species, 121 sites), contained the largest absolute number of significant co-occurrences observed across both taxa. These contrasts provide a varied test bed for asking whether network-selected indicator species can reconstruct the rest of the assemblage.

### 2.2 Presence percentage and reconstruction

Reconstruction accuracy increased with species prevalence for AUC-PR in essentially every cluster and for both taxa (Fig. S3), while AUC remained comparatively flat across the prevalence gradient. This pattern is an expected property of precision-based scoring under class imbalance: for a rare species, even a well-calibrated model faces a much smaller pool of true positives to rank correctly, which mechanically caps achievable precision regardless of the indicator-selection method used, whereas AUC (which conditions on the numbers of positives and negatives separately) is far less sensitive to prevalence. The pattern in Fig. S3 should therefore be read primarily as a property of the evaluation metric rather than as evidence for or against any particular reconstruction method.

### 2.3 Rare species reconstruction

We defined rare species, for this analysis, as those occurring in fewer than 5% of sites among the species retained in the modeling pool; species occurring at fewer than 1% of sites within a cluster were excluded from both the candidate indicator pool and the response-species pool entirely, upstream of any modeling step (see Methods). Because this 1% floor already removes the rarest species, the results below describe reconstruction performance for the less common species that remain, not for the genuinely rarest (< 1%) species. Under this definition, six of the seven bird clusters and one of the four beetle clusters (Cluster 3) contained enough rare species to support a stable estimate; the remaining clusters are omitted from Fig. 3 rather than shown as an uninformative, near-empty panel.

Where rare species could be evaluated, the Consensus method was, on average, at or near the top of the four compared methods on both AUC and AUC-PR in most bird clusters, though confidence bands for the four methods overlapped substantially, particularly at small indicator-set sizes and in clusters with few qualifying species (Fig. 3). Because AUC’s baseline for an uninformative classifier is 0.5 regardless of species prevalence – unlike AUC-PR, whose baseline tracks prevalence itself – the AUC values of roughly 0.6–0.7 observed for rare species across most clusters indicate genuine, non-trivial discriminative skill rather than an artifact of class imbalance. In Beetles Cluster 3, Consensus showed a visible AUC-PR advantage over IndVal, Richness and Random at indicator-set sizes of six and above, while Richness performance was markedly unstable across indicator-set sizes, including AUC values below 0.5 (worse than chance) at some settings. We treat all rare-species results as exploratory, given the small sample sizes per panel even after pooling across indicator-set sizes. Whatever advantage the Consensus method shows here is modest and cluster-dependent, and broadly consistent with its comparable, rather than dramatically superior, performance across all species (Fig. 2).

**Figure 2.**
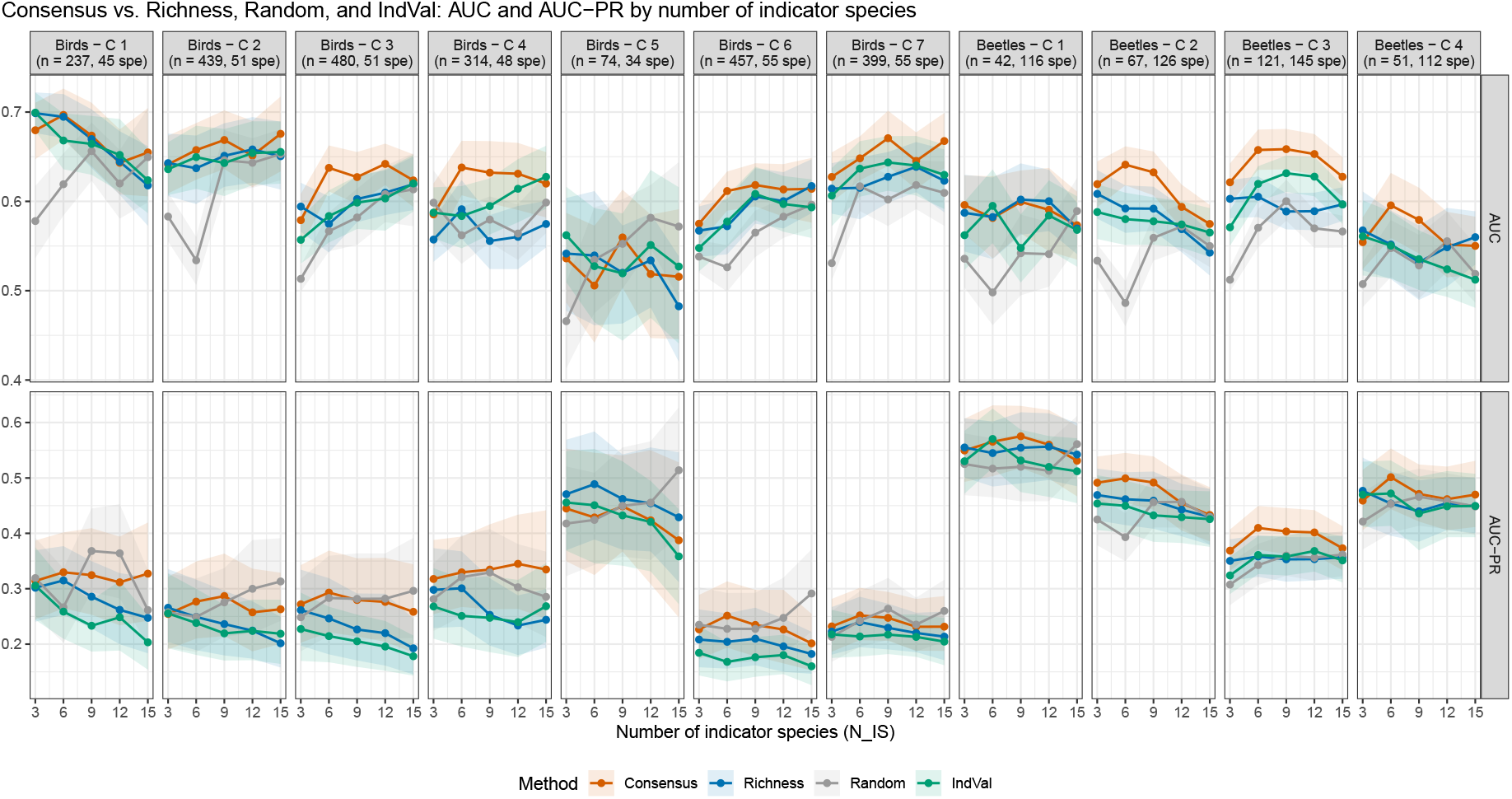
Consensus indicator selection compared against Richness, Random, and IndVal baselines, evaluated exclusively on cross-validated, held-out predictions. Top row: AUC; bottom row: AUC-PR (average precision), both as a function of the number of indicator species (*N*_IS_ = 3, 6, 9, 12, 15), shown separately for every bird cluster (1–7) and beetle cluster (1–4). Points are across-species means; shaded bands are 95% confidence intervals. Site and species counts per panel are given in the panel labels.

**Figure 3.**
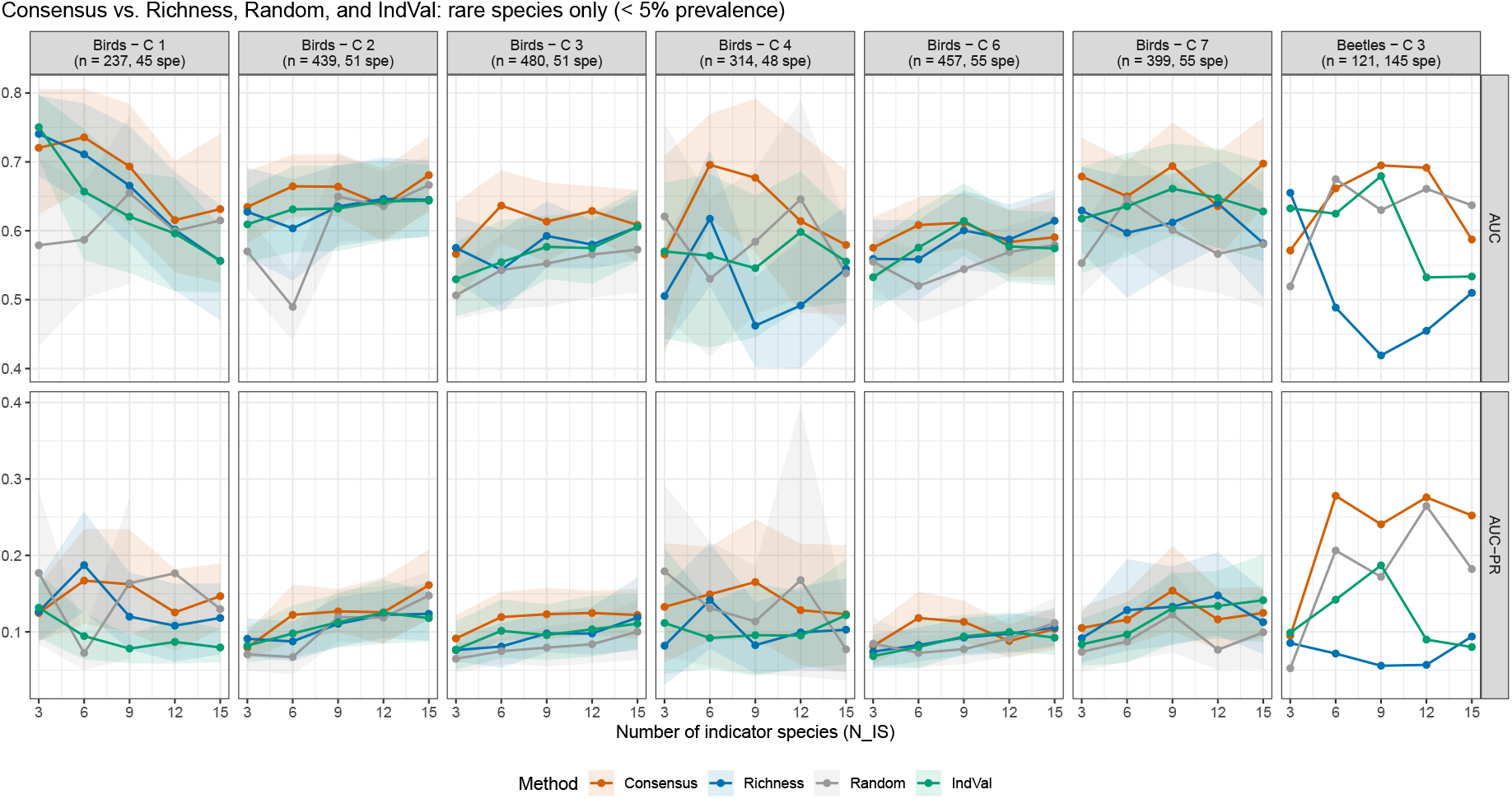
Reconstruction performance for rare species (< 5% prevalence among species retained in the modeling pool) across the Consensus, Richness, Random, and IndVal methods, evaluated exclusively on cross-validated, held-out predictions. Top: AUC; bottom: AUC-PR, as a function of the number of indicator species. Only cluster *×* taxon combinations with at least one qualifying rare species are shown; panels are omitted, not drawn blank, where none exist. Shaded bands are bootstrap 95% confidence intervals, omitted below four qualifying species.

### 2.4 Indicator species versus environmental predictors

For every species, cluster, method, and indicator-set size, we additionally fit occurrence models using climate predictors alone (*Environment*), the selected indicator species alone (*IS*), and their combination (*Environment+IS* ; see Methods), to test whether indicator-based prediction adds information beyond measured climate. This comparison, presented here as an additional analysis after the primary indicator-selection results above, produced a genuinely mixed, assemblage-dependent answer rather than a uniform yes or no (Fig. 4). In Beetles Clusters 1, 2, and 4, IS-alone and Environment+IS scores sat within, or below, the environment-only band for all four headline methods (Consensus, Richness, Random, and IndVal), meaning that indicators performed about as well as climate but added little on top of it. Beetles Cluster 3 was the exception: Consensus and IndVal exceeded the environment-only band, while Richness and Random tracked it more closely, suggesting that a more informative selection criterion can extract signal that the naive baselines do not. Across the bird clusters, indicator-based models (IS alone or Environment+IS) generally sat at or below the environment-only band for all four methods, with the partial exception of the smallest assemblage (Cluster 5, 74 sites), where wide, overlapping confidence bands make any comparison unreliable at this sample size.

**Figure 4.**
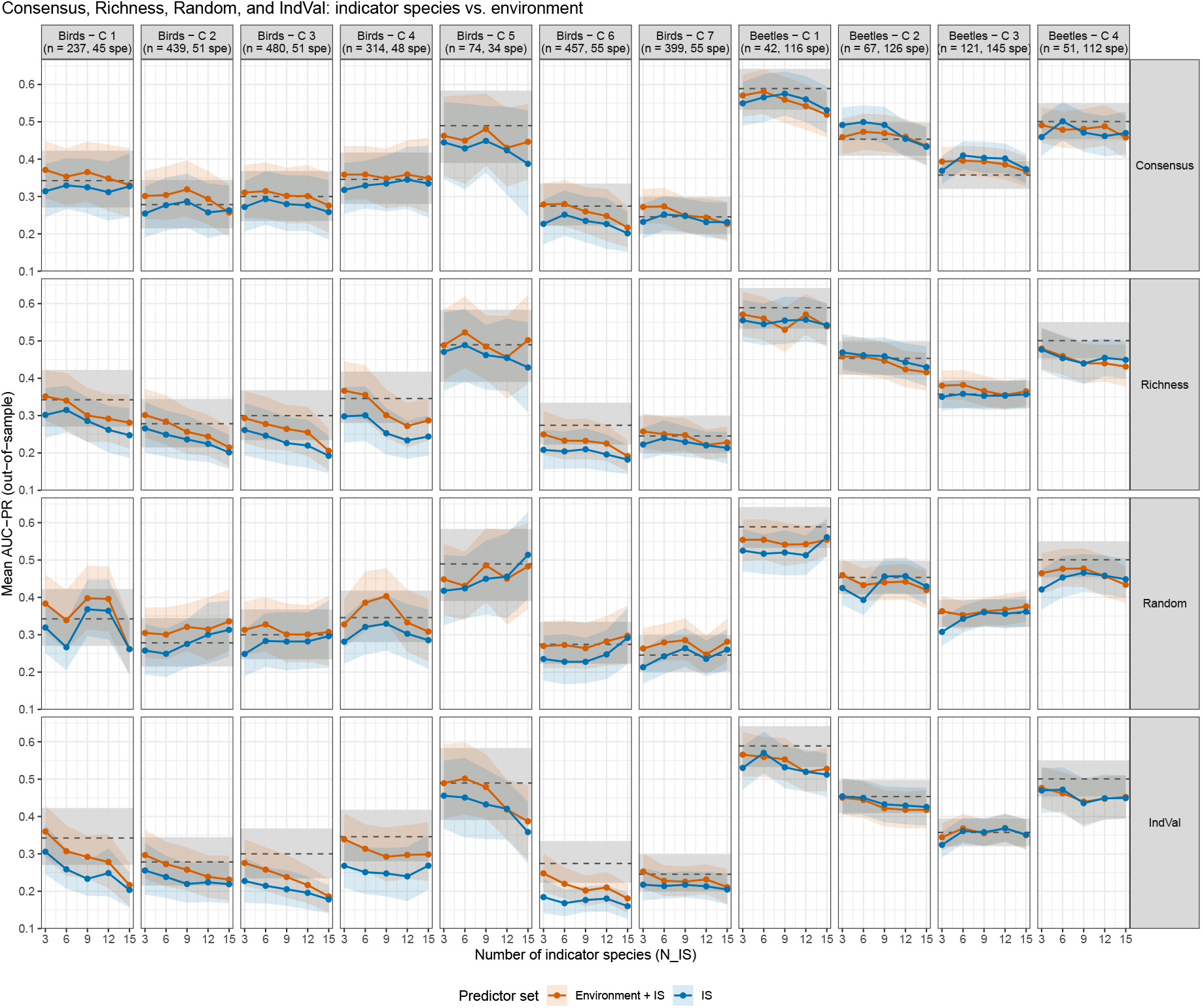
Indicator species versus environmental predictors, evaluated exclusively on cross-validated. Mean out-of-sample AUC-PR (points, with 95% confidence bands) for the IS-only and Environment+IS predictor sets, as a function of the number of indicator species, for the Consensus, Richness, Random, and IndVal methods (rows), by cluster (columns). The dashed line and grey band show the Environment-only baseline (computed once per species per cluster, independent of indicator-selection method and *N*_IS_) and its 95% confidence band.

Read together, these results suggest that indicator species meaningfully substituted for, or added to, environmental information in only one assemblage (Beetles Cluster 3, and there only under the Consensus and IndVal methods), while everywhere else – across all seven bird clusters and the remaining three beetle clusters – measured climate alone predicted community composition about as well as, or better than, indicator-based models built without it. We treat this as an assemblage- and method-specific finding rather than a general claim about indicator species.

Reconstruction performance for beetles was, across these comparisons, at least as strong as for birds despite a community that differs markedly in life-history traits, dispersal ability, and network structure (sparser networks despite far more species; Fig. S2), and the clearest evidence anywhere in this study that indicator species carry information beyond climate came from a beetle assemblage (Cluster 3; Fig. 4). Overall, these results support the transferability of network-guided indicator selection across taxonomically and structurally distinct communities, while underscoring that neither the size of the advantage over baselines nor the added value of indicators over climate is uniform across assemblages or taxa.

## 3 Discussion

The central question of this study was whether a small set of species selected from co-occurrence network structure can provide useful information about the occurrence of the remaining assemblage. Across bird and beetle communities, the Consensus network-based criterion was generally competitive with Richness and IndVal and showed clearer advantages over Random in several assemblages, although the magnitude of these differences varied among assemblages and indicator-set sizes. More importantly, indicator species rarely improved prediction beyond measured environmental variables. These results suggest that species occupying informative positions in a co-occurrence network can be useful for assemblage reconstruction, but structural importance does not necessarily translate into additional predictive information about other species.

The Consensus criterion was designed to integrate complementary network roles rather than rely on a single definition of importance. Positive and negative degree identify species involved in many signed associations, whereas betweenness and BC capture species occupying bridging or less redundant positions in the network. Combining these roles therefore provides a reproducible way to prioritize candidate indicators across different forms of network structure. The fact that Consensus was competitive with conventional approaches, while not uniformly superior to Richness or IndVal, is consistent with the broader literature showing that indicator performance can vary substantially across ecological contexts and environmental gradients (Blanchet et al., 2020; Terrigeol et al., 2022). The contribution of network topology is therefore not that it guarantees the best indicator set in every assemblage, but that it provides an explicit structural basis for selecting species when only a limited subset can be monitored.

The distinction between structural and predictive importance is also consistent with the fact that species are embedded in complex networks of dependencies. Pairwise associations cannot necessarily be interpreted independently of the remaining community, because the apparent relationship between two species may change once their associations with other species are considered (Cazelles et al., 2016; Blanchet et al., 2020). In particular, Cazelles et al. Cazelles et al. (2016) showed that species with higher degree can exhibit weaker statistical association with each of their individual partners, because their occurrence is distributed across a larger number of dependencies. This provides a possible explanation for why species occupying highly connected or central positions in our co-occurrence networks were not necessarily the most informative predictors of individual non-indicator species. Structural importance therefore describes how a species is embedded in the network as a whole, whereas predictive importance is target-specific and depends on how much information that species contributes for a particular response.

The comparison with environmental models provides the clearest interpretation of what these indicators capture. In most assemblages, Environment-only models performed as well as, or better than, models based on IS, and adding IS to environmental predictors yielded little additional gain. Beetles Cluster 3 was the clearest exception, where Consensus and IndVal-selected species exceeded the environment-only baseline. This suggests that, while selected indicators often act primarily as biological proxies for environmental gradients already represented by measured climate variables, they may in some assemblages capture additional ecological information. One possible explanation for the distinct behavior of Beetles Cluster 3 is that it combines the largest number of sampled sites, the highest species richness among the beetle assemblages, and the largest absolute number of significant co-occurrence associations observed across all clusters. Interestingly, this pattern appears more closely related to the amount of significant association structure available in the network than to network density itself, since the densest networks occurred mainly in bird assemblages that did not consistently show the highest reconstruction performance. Its broader spatial extent may further encompass greater habitat heterogeneity, including forests with contrasting disturbance histories. Together, these features may provide a richer and more strongly resolved association structure while also capturing compositional variation not fully represented by the climate variables used in the Environment models (Le Borgne et al., 2018). Under such conditions, species co-occurrence structure may retain information about unmeasured habitat variation and community organization, allowing selected indicator species to contribute predictive information beyond climate alone. The stronger performance observed in beetles Cluster 3 may therefore reflect a combination of greater association information, species richness, sampling extent, and ecological heterogeneity. These explanations remain hypotheses, however, because their relative contributions were not explicitly tested.

Such redundancy is expected because co-occurrence patterns may arise from shared environmental responses, indirect associations, dispersal processes, or direct ecological relationships. Co-occurrence should therefore not be interpreted as direct evidence of ecological interaction (Blanchet et al., 2020; Cazelles et al., 2016; Harris, 2016). In this study, the network is used in a weaker statistical sense: as a representation of assemblage-level association from which candidate indicator species can be selected. The relevant question is thus not whether individual edges represent causal interactions, but whether the association structure they encode can be summarized by a small number of species that are useful for prediction. Previous work has shown that residual association structure can improve species prediction in joint models (Tikhonov et al., 2017; Warton et al., 2015), but our results indicate that this does not imply that structurally prominent species will necessarily provide additional predictive information once environment is already included.

Species prevalence influenced reconstruction performance, particularly for AUC-PR, which increased strongly with prevalence, whereas ROC AUC was comparatively stable. This contrast is consistent with the sensitivity of precision–recall metrics to class imbalance (Saito and Rehmsmeier, 2015). Importantly, however, the framework retained non-random discriminatory ability for several low-prevalence species: AUC values around 0.6–0.7 indicate that their occurrences could still be reconstructed better than chance. This suggests that indicator-based reconstruction remains informative for some uncommon species, even when precision-based performance is constrained by low prevalence. Nevertheless, uncertainty was substantial in clusters containing few rare species, and species occurring at fewer than 1% of sites were excluded from the modeling pool. We therefore interpret the rare-species results as encouraging but exploratory rather than as evidence that the framework resolves the broader challenge of predicting the rarest taxa (Breiner et al., 2015).

Despite marked differences in species richness and network structure between birds and beetles, the same broad pattern emerged across both taxa: network-guided indicator selection can be useful, but neither its advantage over conventional selection methods nor its added value beyond environment is universal. Our results therefore refine, rather than reject, the use of indicator species for biodiversity monitoring. Network structure offers a transparent way to identify compact and interpretable subsets of species, but their predictive value should be evaluated explicitly against environmental baselines rather than inferred from structural importance alone.

## 4 Methods

### 4.1 Empirical data

We investigated assemblage occurrence reconstruction for a bird community network in the Côte-Nord region of Québec, Canada (48°N to 53°N, 65°W to 71°W), spanning 114,118 km^2^. Presence–absence data were collected during the breeding season (late May to mid-July) from 2010 to 2018 as part of the Atlas of Breeding Birds of Québec (Atlas des oiseaux nicheurs du Québec, 2018), using unlimited-distance 5-minute point counts (Bibby et al., 2000). Climate predictors were generated at 250-m resolution using the BioSIM platform (Régnière et al., 2017), which simulates daily meteorological variables including temperature, precipitation, humidity, and wind from spatially referenced weather-station data. Climate estimates were spatially interpolated using spatial regression and kriging, with elevation as a drifting variable, to account for geographic variation in latitude, longitude, and elevation (Boulanger et al., 2018). A total of 15,000 georeferenced points across Québec were used to generate the high-resolution climate surfaces (see Table 1 in the Supporting Information for the complete list of variables).

To further validate the framework, we incorporated an independent beetle assemblage compiled from published datasets collected in 2004, 2005, 2007, and 2011, comprising 146 beetle species (Janssen et al., 2009; Légaré et al., 2011; Bichet et al., 2016; Bouderbala et al., 2023b). We additionally included 54 sites sampled in 2018 in the northern part of the study region, along the main northeastern road toward Labrador. Beetle sampling involved one multidirectional flight-interception trap per site to capture flying beetles and four meshed pitfall traps per site to sample beetles moving on the soil surface (Janssen et al., 2009; Bichet et al., 2016).

### 4.2 Assemblage identification

Static IS are unlikely to perform consistently over broad climatic extents (Morelli, 2015; Terrigeol et al., 2022). We therefore defined assemblages according to climatic gradients to account for environmental variation in IS composition. Climate clusters were identified separately for birds and beetles using hierarchical clustering on principal components implemented in `FactoMineR` (Husson et al., 2013). Clustering was based on the first two principal components of the climate variables, which together explained more than 95% of the variance.

### 4.3 Co-occurrence networks

Species co-occurrence networks were constructed within each climate cluster from a binary community matrix (species *×* sites), where 1 indicated presence and 0 absence. Nodes represented species and edges represented statistically significant deviations from random co-occurrence. Networks were signed but unweighted: positive edges indicated aggregation, where co-occurrence was more frequent than expected under independence, whereas negative edges indicated segregation, where co-occurrence was less frequent than expected. Summary statistics for each cluster and taxon, including the number of species retained, the number and sign of significant co-occurrence links, and network density, are reported in Fig. S2.

To assess pairwise species associations, we applied the probabilistic co-occurrence model of Veech Veech (2013), implemented in the `cooccur` package (Griffith et al., 2016). Unlike randomization-based null models, this approach uses the hypergeometric distribution to estimate the probability that two species co-occur at exactly *k* sites given their individual frequencies. The probability is

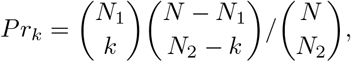

where *N* is the total number of sites and *N*_1_ and *N*_2_ are the numbers of sites occupied by the two species. The observed number of co-occurrences, *Q*_obs_, was compared with this distribution using one-tailed probabilities. The lower-tail probability 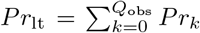 quantified significant segregation, whereas the upper-tail probability 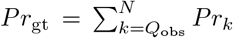 quantified significant aggregation. An edge was added when either *Pr*_lt_ *< α* (negative edge) or *Pr*_gt_ *< α* (positive edge), with *α* = 0.05. Species pairs showing no significant deviation were left unconnected.

### 4.4 Network-based and baseline IS identification

Each climate cluster was represented by a set of *N*_IS_ indicator species selected from a larger pool of *q* potential indicator species. We considered five network-based selection criteria. The first criterion, **Positive**, selected the *N*_IS_ species with the largest number of significant positive co-occurrence links. The second, **Negative**, selected the species with the largest number of significant negative links. The third criterion, **Betweenness**, ranked species according to normalized betweenness centrality,

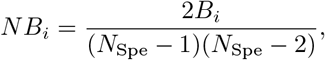

where 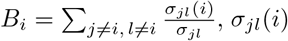 is the number of shortest paths between species *j* and *l* passing through *i*, and *σ*_*jl*_ is the total number of shortest paths between *j* and *l*. Betweenness was calculated on the unsigned co-occurrence network using the `igraph` package (Csardi and Nepusz, 2006). The fourth criterion, **BC**, first identified the set *Spe*_*B,q*_ containing the *q* species with the highest betweenness scores. From this subset, the *N*_IS_ most heterogeneous species were selected using normalized closeness centrality,

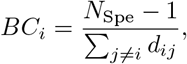

where *d*_*ij*_ is the shortest-path distance between species *i* and *j*, and *N*_Spe_ is the total number of species in the corresponding co-occurrence network. The *N*_IS_ species with the lowest *BC*_*i*_ values were retained, thereby reducing redundancy by avoiding the selection of species that are highly connected and likely to occupy similar positions in the network.

The fifth criterion, **Consensus**, combined information from Positive, Negative, Betweenness, and BC. For a given cluster and *N*_IS_, each base method produced a top-*N*_IS_ ranked list. Each species received one vote for each list in which it appeared, together with a within-list score decreasing from 1 for the top-ranked species to approximately 1*/L* for the lowest-ranked species in a list of length *L*. Species absent from a list received neither a vote nor a score from that method. Species were ranked first by total votes and then by summed within-list score, with alphabetical order used to break remaining ties. The top *N*_IS_ species formed the Consensus indicator set. This procedure favors species that perform consistently across several network roles rather than species that rank highly according to only one structural criterion.

In addition to the five network-based criteria, we considered three baseline methods. The **Richness** baseline selected the *N*_IS_ species whose combined presence pattern best tracked total site-level species richness, following common practice in indicator-species studies that use aggregate richness as a monitoring target (Terrigeol et al., 2022). The **Random** baseline selected *N*_IS_ species uniformly at random from the full species pool independently within each cross-validation fold, providing a null reference.

The **IndVal** baseline used the classic Dufrêne–Legendre indicator value (Dufrêne and Legendre, 1997). Because this method requires sites to be partitioned into ecologically meaningful groups *a priori*, and no independent habitat classification was available, sites were divided into terciles of site-level species richness (low, medium, and high richness), each containing approximately one third of the sites. This allowed IndVal to identify species associated with the richest sites rather than with predefined habitat classes.

For species *i* and site group *j*, the indicator value combines specificity and fidelity,

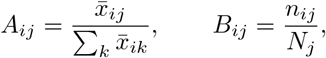

where 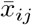 is the mean abundance, or occurrence frequency for presence–absence data, of species *i* in group *j*; the denominator of *A*_*ij*_ sums across all site groups *k*; *n*_*ij*_ is the number of sites in group *j* where species *i* is present; and *N*_*j*_ is the total number of sites in group *j*. Specificity, *A*_*ij*_, is h ighest when o ccurrences are concentrated in group *j*, whereas fidelity, *B*_*ij*_, i s h ighest w hen s pecies *i* o ccurs a t m ost s ites i n t hat group. The indicator value is

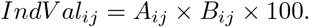

Statistical significance was assessed using 999 permutations. Site labels were randomly reassigned among the three richness groups while preserving group sizes and species occurrence records, and *IndV al*_*ij*_ was recalculated for each permutation to generate a null distribution. The *p*-value was defined as the proportion of permuted values at least as large as the observed value. For each cluster, the *N*_IS_ species with the highest significant (*p* ≤ 0 .05) IndVal scores f or the h igh-richness g roup were selected.

### 4.5 Species occurrence modeling

To infer the occurrence probability of each non-indicator species (non-IS), we fitted binomial generalized linear models (GLMs) to presence–absence observations. Species distribution models combine occurrence observations with environmental predictors to estimate and predict species distributions (Elith and Leathwick, 2009); here, we used the same occurrence-modeling framework to compare the predictive information carried by selected indicator species and by measured climate variables.

For every non-IS species within each climate-defined assemblage, we fitted three predictor sets: selected indicator species alone (**IS**); climate predictors alone (**Environment**: mean, maximum, and minimum temperature, and precipitation); and both sets jointly (**Environment+IS**). This design distinguishes whether indicator species can substitute for measured climate from whether they provide complementary predictive information beyond climate. The Environment-only model was fitted once per species per cluster, independently of indicator-selection method and *N*_IS_, because its predictors do not depend on the selected indicator set.

For each non-IS species *j* within a climate cluster, the models estimated the conditional probability of occurrence at site *S*_*n*_ given the corresponding indicator-species and/or climate predictors,

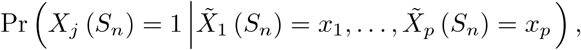

where *x*_*i*_ denotes either the occurrence of an indicator species (∈{0, 1}) or a continuous climate variable at site *S*_*n*_, and *p* is the number of predictors in the relevant predictor set.

Model performance was evaluated from out-of-fold predictions obtained by five-fold cross-validation. For each species, models were fitted on four folds and evaluated on the held-out fold, so every site contributed once to the evaluation set. We report ROC AUC and AUC-PR (average precision). AUC-PR complements ROC AUC for low-prevalence species because it is less driven by correctly classified absences and more directly reflects the precision–recall trade-off for rare binary events.

## Code Availability

Code supporting the findings of this study, including the full cross-validated analysis pipeline described above, will be made available in a public repository at the time of resubmission, together with a permanent archival DOI.

## Acknowledgements

This work was supported by the Sentinel North program of Laval University, funded by the Canada First Research Excellence Fund. AA, DF and PD were also supported by the Natural Sciences and Engineering Research Council of Canada (NSERC). We acknowledge Calcul Québec and Compute Canada for their technical support and computing infrastructures. We thank also the Québec Breeding Bird Atlas for supplying data. We would also like to thank the following partners: Regroupement QuébecOiseaux, Environment and Climate Change Canada and Birds Canada, together with all the volunteer participants who gathered data for the project. We are grateful to Louis-Paul Rivest for his valuable statistical suggestions and advice.

## Notes

### Competing Interest Statement

The authors have declared no competing interest.

### Summary of Updates

This version includes the following main revisions. 1. Strengthened statistical validation. Model performance is now evaluated strictly out of sample using cross-validation, with model and threshold selection separated from evaluation. This provides more conservative and robust performance estimates than in the previous version. 2. Refined interpretation of rare species results. Under the revised validation procedure, performance for rare species is lower than previously reported. Some low-prevalence species are still predicted better than chance, and we now interpret these results more cautiously. 3. New primary performance metrics. The Sorensen-Dice dissimilarity computed with a fixed threshold of 0.5 is no longer the main measure. It was replaced by two threshold-independent metrics, AUC and AUC-PR. AUC-PR is especially informative for rare species because its baseline depends on species prevalence, unlike AUC, whose baseline remains 0.5, providing a more demanding assessment under strong class imbalance. 4. Additional benchmark method. We added the classical Dufrene-Legendre indicator value index, IndVal, as a third benchmark alongside the Richness and Random approaches. 5. Updated results and conclusions. All numerical values and main conclusions were rechecked against the updated figures and results, including network density, rare species, and the comparison between indicators and environmental variables. Several statements were revised to better reflect the updated results. 6. Explicit comparison with environmental variables. We added a comparison of models using only environmental variables, only indicator species, and both combined, to test whether indicator species provide information beyond measured climate or mainly act as a biological proxy for it.

