## Supplementary material for "When indicator species identified from a network structure predict biodiversity": SI

**This PDF file includes:**

Figures S1 to S5

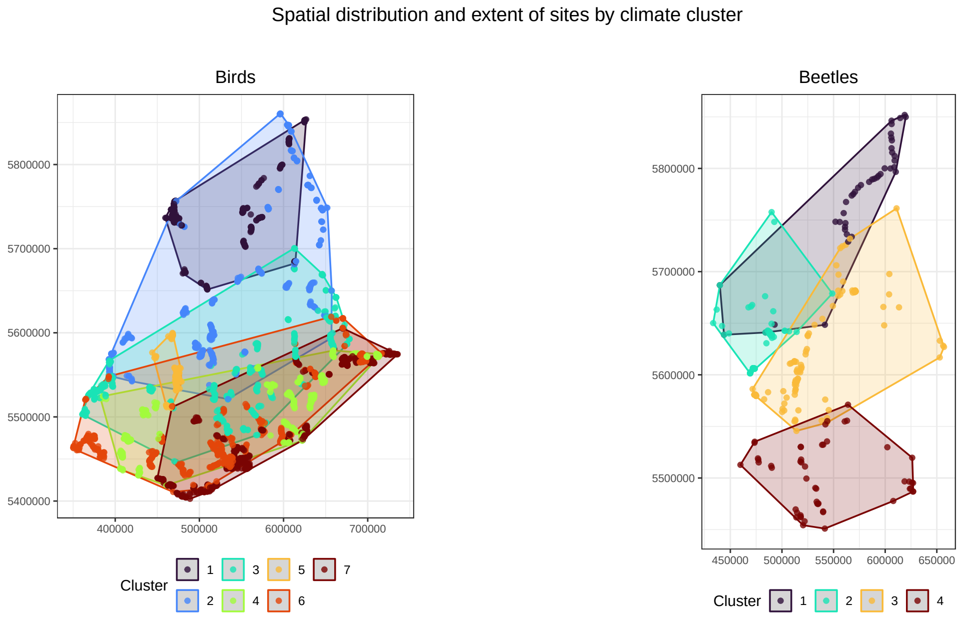


**Figure S1.** Spatial distribution of sampling sites by climate-defined cluster, shown separately for birds (seven clusters, fully automatic clustering, all retained) and beetles (four clusters, fixed cluster count). Coordinates are the projected site coordinates (in meters) used throughout the analysis. Each cluster’s spatial extent is additionally shown as a 100% minimum convex polygon (MCP).

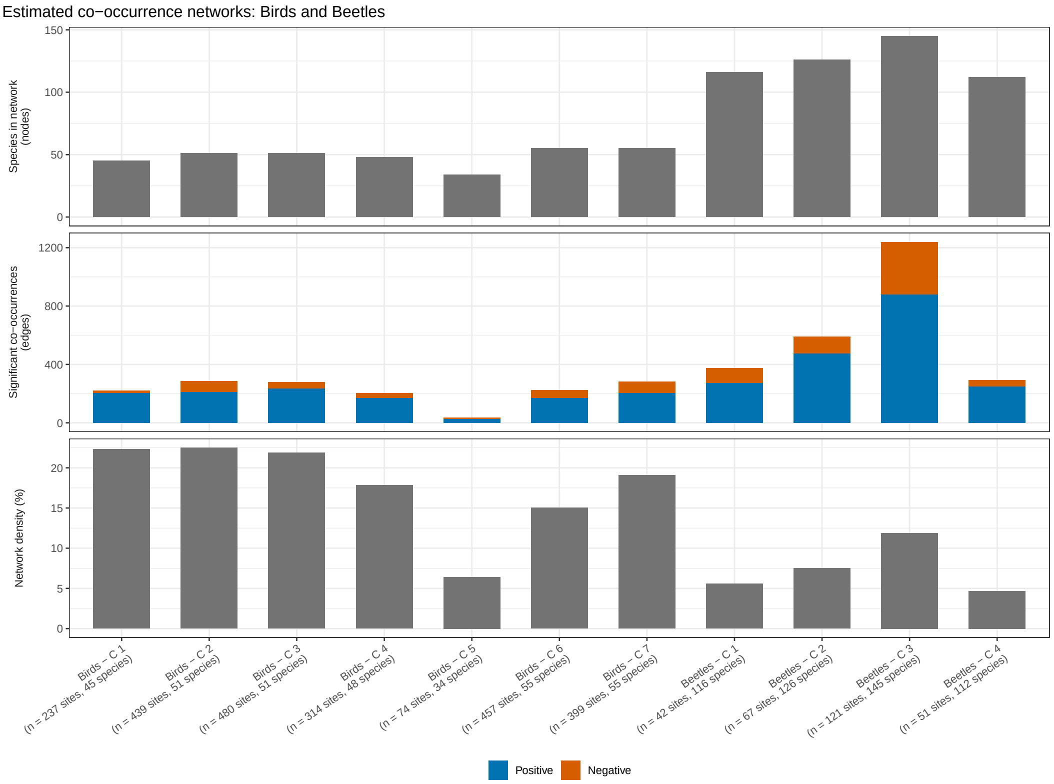


**Figure S2.** Structure of the estimated co-occurrence networks for birds (seven clusters) and beetles (four clusters). Top: number of species retained in the network (nodes). Middle: number of statistically significant co-occurrences (edges), split into positive and negative associations. Bottom: network density, the percentage of all possible species pairs connected by a significant edge. Site and species counts per cluster are given in the panel labels.

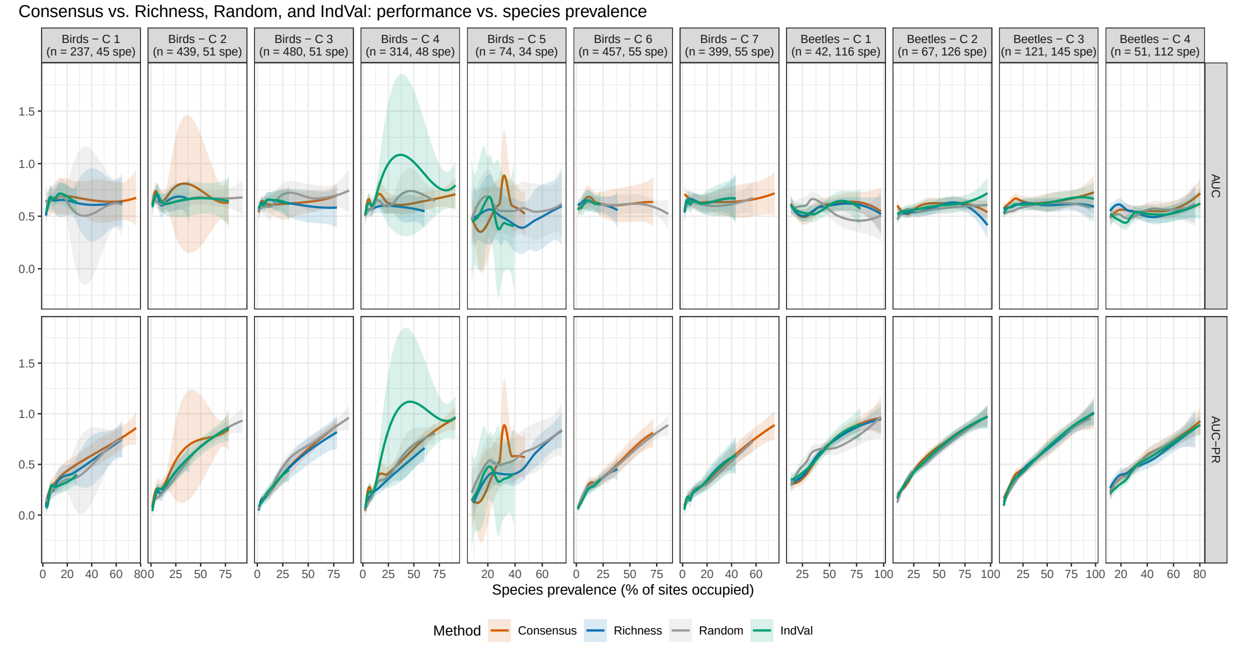


**Figure S3.** Relationship between species prevalence and reconstruction performance, evaluated exclusively on cross-validated, held-out predictions. AUC (top) and AUC-PR (bottom) as a function of species prevalence (% of sites occupied), by cluster (Birds 1–7, Beetles 1–4), for the Consensus, Richness, Random, and IndVal methods (loess curves with 95% confidence bands, pooled across indicator-set sizes). Higher values indicate better performance.

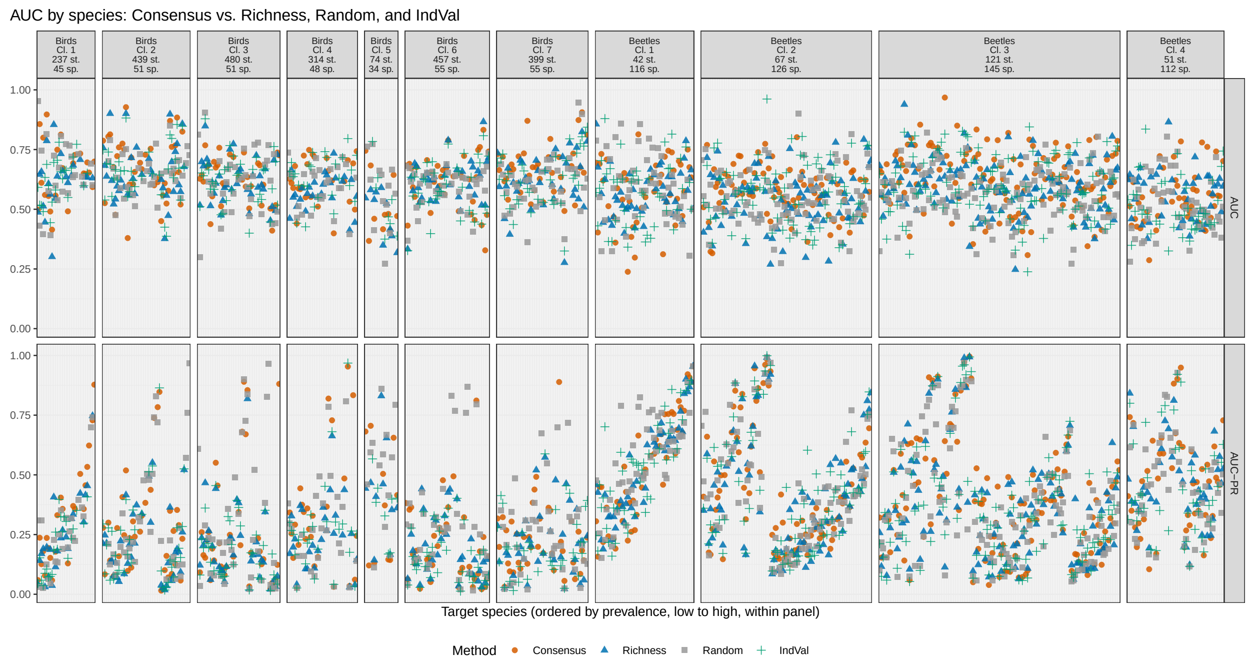


**Figure S4.** Species-level reconstruction performance for the four headline indicator-selection methods. AUC and AUC-PR are shown for individual target species within each bird and beetle assemblage. Species are ordered by prevalence within each panel.

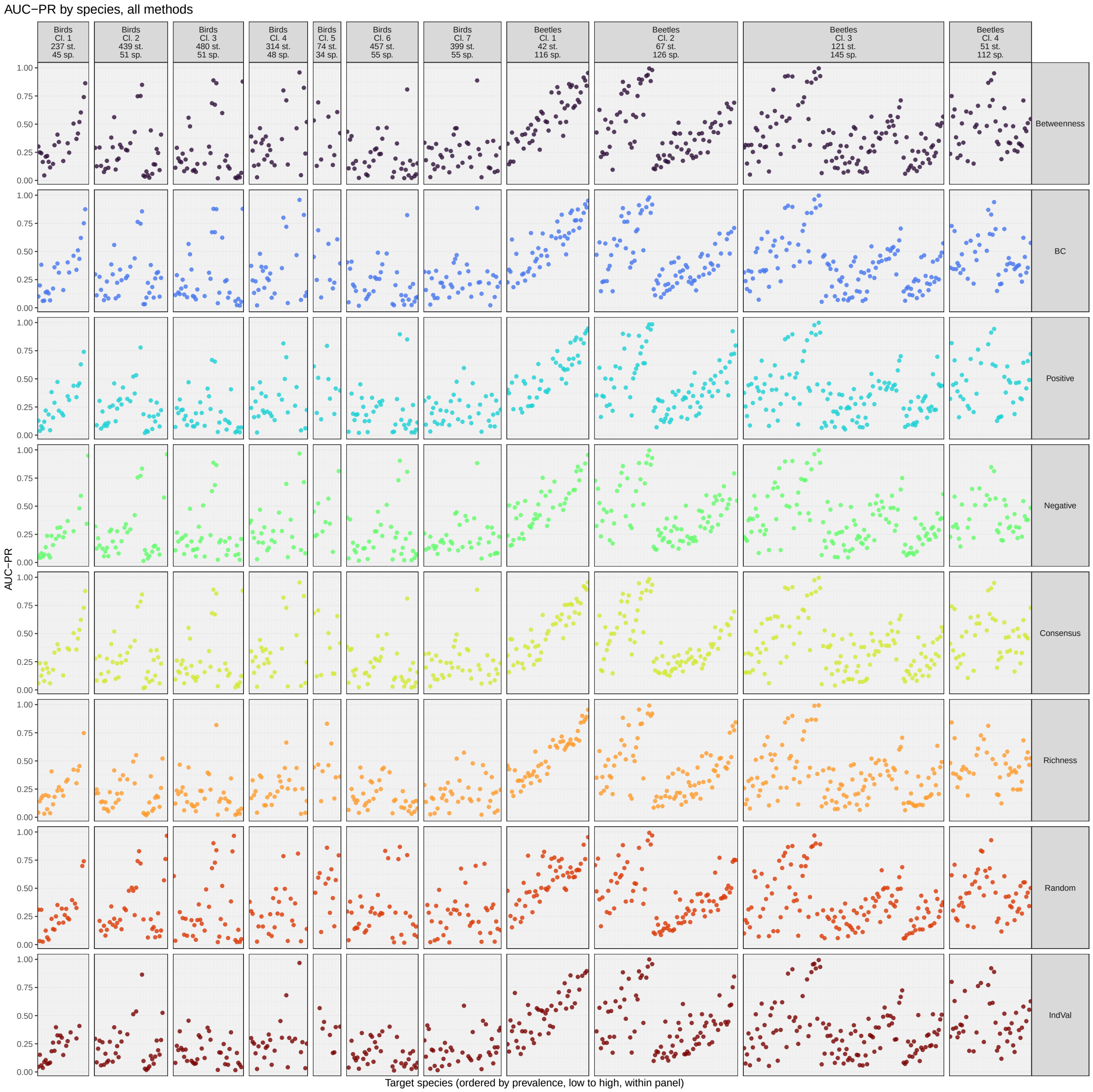


**Figure S5.** Species-level AUC-PR across all indicator-selection methods. Results are shown for individual target species within each bird and beetle assemblage, with species ordered by prevalence within each panel.
